# Single cell transcriptional and functional analysis of human dopamine neurons in 3D fetal ventral midbrain organoid like cultures

**DOI:** 10.1101/2020.10.01.322495

**Authors:** Marcella Birtele, Yogita Sharma, Petter Storm, Janko Kajtez, Jenny Nelander Wahlestedt, Edoardo Sozzi, Fredrik Nilsson, Simon Stott, Xiaoling L He, Bengt Mattsson, Daniella Rylander Ottosson, Roger A Barker, Alessandro Fiorenzano, Malin Parmar

## Abstract

Transplantation of midbrain dopamine (DA) neurons for the treatment of Parkinson’s disease (PD) is a strategy that has being extensively explored and clinical trials using fetal and stem cell-derived DA neurons are ongoing. An increased understanding of the mechanisms promoting the generation of distinct subtypes of midbrain DA during normal development will be essential for guiding future efforts to precisely generate molecularly defined and subtype specific DA neurons from pluripotent stem cells. In this study, we used droplet-based scRNA-seq to transcriptionally profile a large number of fetal cells from human embryos at different stages of ventral midbrain (VM) development (6, 8, and 11 weeks post conception). This revealed that the emergence of transcriptionally distinct cellular populations was evident already at these early timepoints. To study late events of human DA differentiation and functional maturation, we established a primary fetal 3D culture system that recapitulates key molecular aspects of late human DA neurogenesis and sustains differentiation and functional maturation of DA neurons in a physiologically relevant cellular context. This approach allowed us to define the molecular identities of distinct human DA progenitors and neurons at single cell resolution and construct developmental trajectories of cell types in the developing fetal VM.

Overall these findings provide a unique transcriptional profile of developing fetal VM and functionally mature human DA neurons, which can be used to quality control stem cell-derived DA neurons and guide stem cell-based therapies and disease modeling approaches in PD.

## INTRODUCTION

DA neurons in the ventral midbrain are diverse and consist of several different subtypes. of these, the A9 DA neurons in substantia nigra compacta (SNc) are selectively lost in PD, whereas nearby A10 DA neurons of the ventral tegmental area (VTA) are relatively spared by the disease [8]. However, little is known about how these different dopaminergic populations of cells develop and recent scRNA-seq studies of adult mouse midbrain have revealed a greater than expected molecular diversity in mature DA neurons suggesting heterogeneity even within anatomically defined DA clusters [9–11]. Given the importance of DA neurons in a broad spectrum of applications such as disease modeling, diagnostics, drug screening as well as in cell-based therapies for PD, major efforts to develop more refined and precise differentiation protocols to generate midbrain DA neurons of precisely defined molecular identities from pluripotent stem cells (PSCs) are continuously on-going, and a better understanding of human DA neuron specification and maturation is vital in these efforts.

Single cell RNA-sequencing (scRNA-seq) represents a major technological advancement in determining cell type and developmental trajectory, as it provides an insight into dynamic gene expression at single cell resolution with potential to analyze cell-cell gene expression variability [10, 12, 13], and scRNA-seq has already increased our understanding of aspects DA neuron development [10, 14, 15]. Due to extremely limited access to human fetal brain tissue, such investigations have almost exclusively been performed in mice [11, 15, 16]. One study used single cell transcriptomics to compare DA lineage in mouse and human development, which revealed several points of similarity but also critical differences between species [10], highlighting the importance of studying midbrain development also using human tissue.

In this study, we used droplet-based scRNA-seq to perform high-throughput transcriptional profiling of a large number of cells from human fetal midbrain at different developmental stages. Our analysis identified the cellular composition of the developing VM during dopaminergic neurogenesis, and revealed the early emergence of transcriptionally distinct DA populations. To further study late stages of human DA neurogenesis and to investigate molecular diversity in human DA neurons once they have become functionally mature, we used primary cultures of human fetal VM. However, we found that standard primary 2D cultures did not maintain human DA neurons for extended periods, and instead we established an organoid-like 3D culture model. Using scRNA-seq transcriptomics, we identified two major molecularly distinct DA progenitors and two major DA neuron subtypes.

## RESULTS

### Single cell transcriptomics reveals cellular composition of the developing human fetal VM

Human DA progenitors located in the ventral midbrain gives rise to post-mitotic DA neurons during early brain development (Fig. 1A) [17, 18]. At mid DA neurogenesis (week 7.5 post conception [PC]), SOX2 marks the proliferative ventricular zone (VZ), OTX2 is expressed from the ventricular to mantle zone, and *TH* expression labels DA post-mitotic neurons (Fig. 1B). iDISCO tissue clearing was used for a 3D anatomical reconstruction of the VM, and through using TH mapping we could identify intricate DA projections to the forebrain target regions already at these early timepoints (week 7–11 PC) (Fig. 1C). To determine the cellular composition of the developing VM at the molecular level, we subdissected VM from human embryos of gestational ages ranging from 6 to 11 weeks PC, spanning early to peak DA-genesis [18–20].

**Figure 1.**
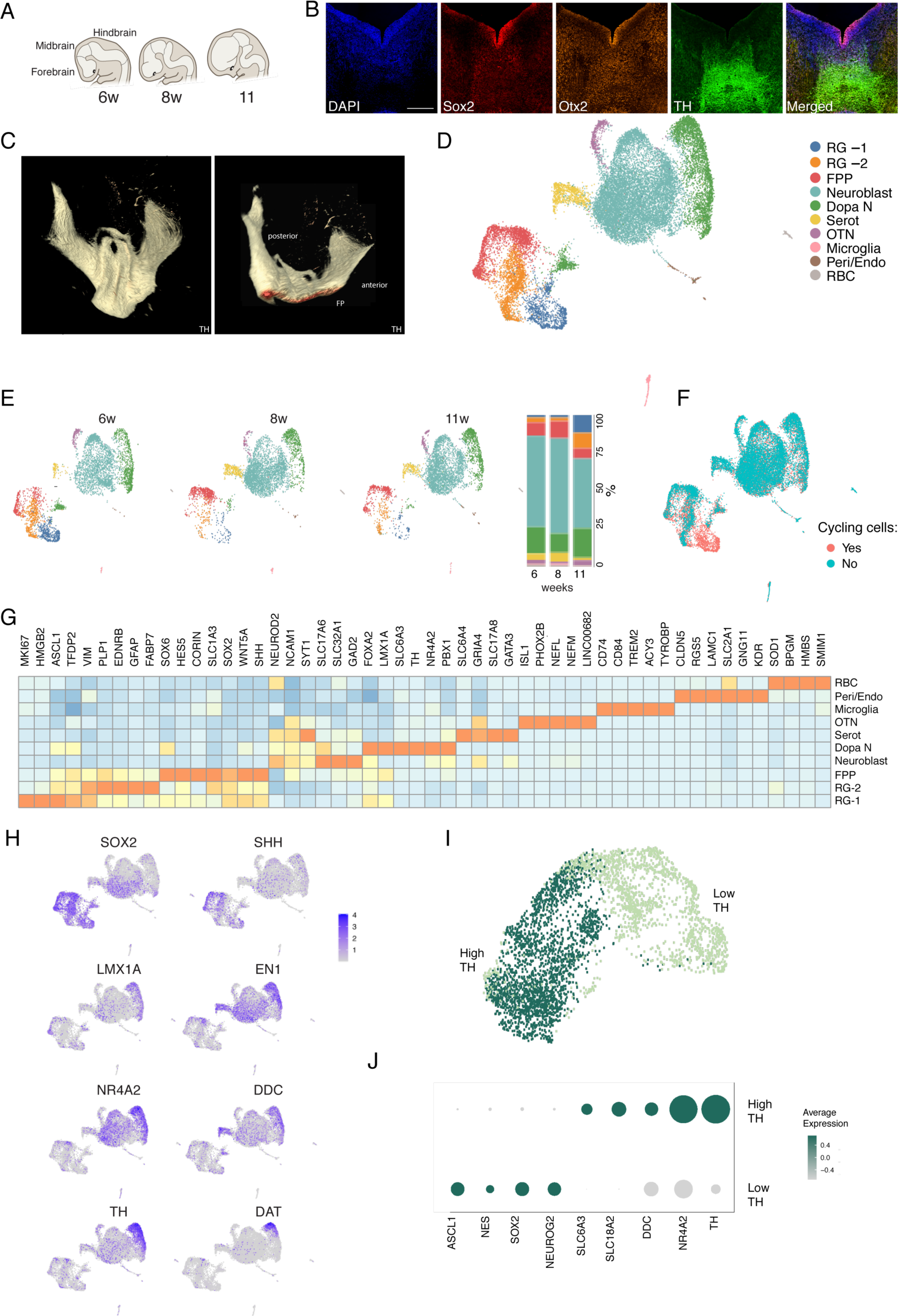
sc-RNAseq of developing human fetal VM. **A**, Schematic representation of the formation of different structures during human brain development (6 to 11 weeks PC). **B**, Immunohistochemistry of SOX2, OTX2 and TH in human VM coronal section at PC week 7.5. Scale bars, 250 µM. **C**, iDISCO circuitry reconstruction obtained by mapping TH in in a human fetal VM at PC week 7.5. **D**, Uniform manifold approximation and projection (UMAP) plot showing clustering of 23,483 analyzed cells from three separate fetuses (6, 8, 11 weeks PC). Cell type assignments are indicated. **E**, UMAP plots of cell clusters for each developmental stage of fetal VM analyzed and the fraction (%) of cells per cluster. **F**, UMAP plots showing cell cycle scores of analyzed cells (S.Score and G2M.Score). Cycling cells showed with blue circles. **G**, Heat map showing expression levels of indicated genes per cluster. Indicated genes are established markers for radial glial, floor plate, DA, Serot, OTN, microglia, Peri/Endo and RBC cells. **H**, Feature plots visualizing early and late DA markers across clusters. Colors indicated expression level. **I**, UMAP plot showing DA subclusters and **j**, dot plot showing differentially expressed selected genes.

We performed droplet-based scRNA-seq on the dissected VM tissue from three separate fetuses (6, 8, 11 weeks PC) (Fig. 1D-E). After removal of low-quality cells (see Materials and Methods), a total of 23,483 cells were retained for analysis. Uniform manifold approximation and projection (UMAP) graph-based clustering partitioned the cells into 10 distinct clusters (Fig. 1D). Three small but clearly distinct clusters were identified as microglia (*CD74, CD86, TREM2, ACY3, TYROBP*; 0–2% of total cell population across all embryos), pericytes and endothelial cells (*CLDN5, RGS5, LAMC1, GNG11, KDR*; 0–2% of total cell population across all embryos), and red blood cells (*SOD1, BPGM, HMBS, SLC2A1, SMIM1*; 0–1% of total cell population across all embryos), which are expected contaminants after fetal tissue dissection (Fig. 1D,G). The majority of cells fell into the remaining seven clusters, which we were able to assign to major cell types through analysis of canonical and unbiased markers.

We found two major clusters sharing key molecular features of radial glial (RG) cells, which we termed RG-1 (blue) and RG-2 (orange). Both of these clusters were characterized by expression of *SOX2, PLP1, EDNRB*, and *SOX9* and they were proportionally reduced during later developmental stages of human VM embryos (Fig. 1D,E and Supplementary Fig. 1C). RG-1 mainly included cycling RG cells in the ventricular zone and were distinguished by a highly proliferative signature (*TOP2A, MIKI67, TFDP2*), as visualized by UMAP cell cycle score (S.Score and G2M.Score), as well as by pro-neural gene *ASCL1* and chromatin-associated gene *HMGB2a* expression (Fig. 1F and Supplementary Fig. 1A,B). RG-2 on the other hand showed higher expression of canonical RG markers, including *FABP7* (also known as BLBP1) and *SLC1A3* (also known as GLAST), linking RG-1 with floor plate progenitors (FPP; red) (Fig. 1D). This latter cluster was enriched by expression of morphogens *SHH* and *WNt5A* as well as *FOXA2, LMX1A*, and *OTX2*, which define midbrain FP cells (Fig. 1D, G and Supplementary Fig. 1C).

A predominant cluster (teal), mainly comprising post-mitotic cells with heterogeneous cell identities, referred to here as neuroblasts (Nb), was also detected, and increased proportionally during fetal VM development (Fig. 1D,G). This cluster was characterized by the loss of proliferative markers (Fig. 1F and Supplementary Fig. 1A,B) and expression of the neuronal differentiation factor 2 (*NEUROD2*), the cytoskeletal marker neural cell adhesion molecule 1 (*NCAM1*), and the synaptic marker *SYT1*. Emerging cell type diversity including expression of genes associated with DA neuron progenitors (*FOXA2, NR4a2 EN2, NR4A2, LMX1A, TH*) and serotoninergic (Serot) progenitors (*GRIA4, GATA3*) was revealed in this cluster (Fig. 1D,G and Supplementary Fig. 1C). These immature neuronal cells subsequently differentiated into three types of mature neurons: Serotonergic neurons (yellow cluster expressing *SLC6A4, GRIA4, SLC17A8, GATA3*; 2–6% of total cell population across all embryos) and oculomotor/trochlear nucleus (OTN) neurons (purple cluster expressing *ISL1, PHOX2B, NEFL, NEFM, LINC00682*; 1–3% of total cell population across all embryos) and a much larger population of DA neurons (green cluster expressing *TH, DDC* and *NR4A2* (also known as *NURR1*) (13–20% of total cell population across all embryos) (Fig. 1D,G). *TH* and DA transporter (*DAT*) expression was not homogeneously expressed in the DA cluster however, prompting us to further investigate gene expression at the single cell level within this population (Fig. 1H and Supplementary Fig. C).

By iterating the resolution parameter on DA clusters from different embryos (Fig. 1H), we distinguished the most mature DA neuron population and identified two subclusters with high and low *TH* expression (named TH^high^ and TH^low^, respectively) (Fig. 1I). TH^low^ was enriched in early DA markers such as *SOX2, EN1*, and *SOX6*, while the TH^high^ population was characterized by higher expression of ion channels as well as by expression of *DAT* and the synaptic vesicular transporter *SLC18A2* (also known as *VMAT2*), denoting a more advanced DA differentiation stage (Fig. 1I,J and Supplementary Fig. 1E).

This data provides a time-resolved single cell map of the developing human fetal VM and shows that emerging molecularly distinct DA neuron can already be identified at this stage of development. Further investigations into fetal VM at later stages of development would be highly desirable, but access to second trimester fetal tissue is very restrictive and cannot be easily used for studies of this type.

Thus in order to undertake transcriptional studies of functionally mature human DA neurons, we established primary cultures from hVM collected from three fetuses (8, 9, 10 weeks PC) (Supplementary Fig. 2A,B). After two weeks of differentiation, a large number of TH+ cells with neuronal morphology were detected in these cultures (Fig. 2A). Whole-cell patch-clamp recordings at day 15 showed that the cells (n=19) had a resting membrane potential (RMP) comparable to a mature neuronal profile (Fig. 2C and Supplementary Fig. 2H,I). In line with these findings,, the cells were able to fire induced action potentials (APs) (Fig. 2D,E). At this timepoint, we found that 23.5% of the cells recorded were also able to spontaneously fire APs (Fig. 2D and Supplementary Fig. 2J), suggesting that they had started a process of functional maturation. To assess more mature DA neurons, we attempted to analyze the neurons after an additional two weeks in culture (Supplementary Fig. 2A,B). However, cells with a neuronal morphology were few and sparsely distributed at this stage (Fig. 2B). Immunocytochemical analysis showed that very few, if any, TH neurons remained in the cultures (Fig. 2B and Supplementary Fig. 2D,G). In line with this finding, when recording cells at day 30, no inward Na^+^ and outward K^+^ voltage-dependent currents, induced APs, or spontaneous firing were observed (n=23 cells) (Fig. 2C-E and Supplementary Fig. 2H-J). The depletion of TH neurons over time in 2D culture was accompanied by expansion of immature neuronal cells (NESTIN, SOX2) (Supplementary Fig. 2C,D) and emergence of non-neuronal populations containing glial (GFAP) and oligodendrocyte (OLIG2) progenitors (Supplementary Fig. 2E,F). To transcriptionally characterize the cell type composition in these primary cultures at these two different timepoints (day 15 and day 30), we generated a single cell dataset of 22 665 QC-filtered cells from two embryos that were grouped into 10 cell types using graph-based clustering (Fig. 2F,G). To assign cluster identities, we used the cell types previously defined in uncultured fetal VM and projected these labels onto the 2D culture dataset. *In silico* analysis identified similar clusters in 2D culture to those in uncultured fetal VM tissue, including RG (*BLBP, GLAST*) and FPP populations (*SHH, FOXA2*) as well as a DA cluster (*NURR1, TH*) (Fig. 2F,G and Supplementary Fig. 2K). Two clusters with low predictions scores and no direct correspondence to any uncultured fetal VM populations were identified as oligodendrocyte (*OLIG1, OLIG2*) and astrocyte (*GFAP, AQP4*) progenitor clusters (Fig. 2F,G and Supplementary Fig. 2K).

**Figure 2.**
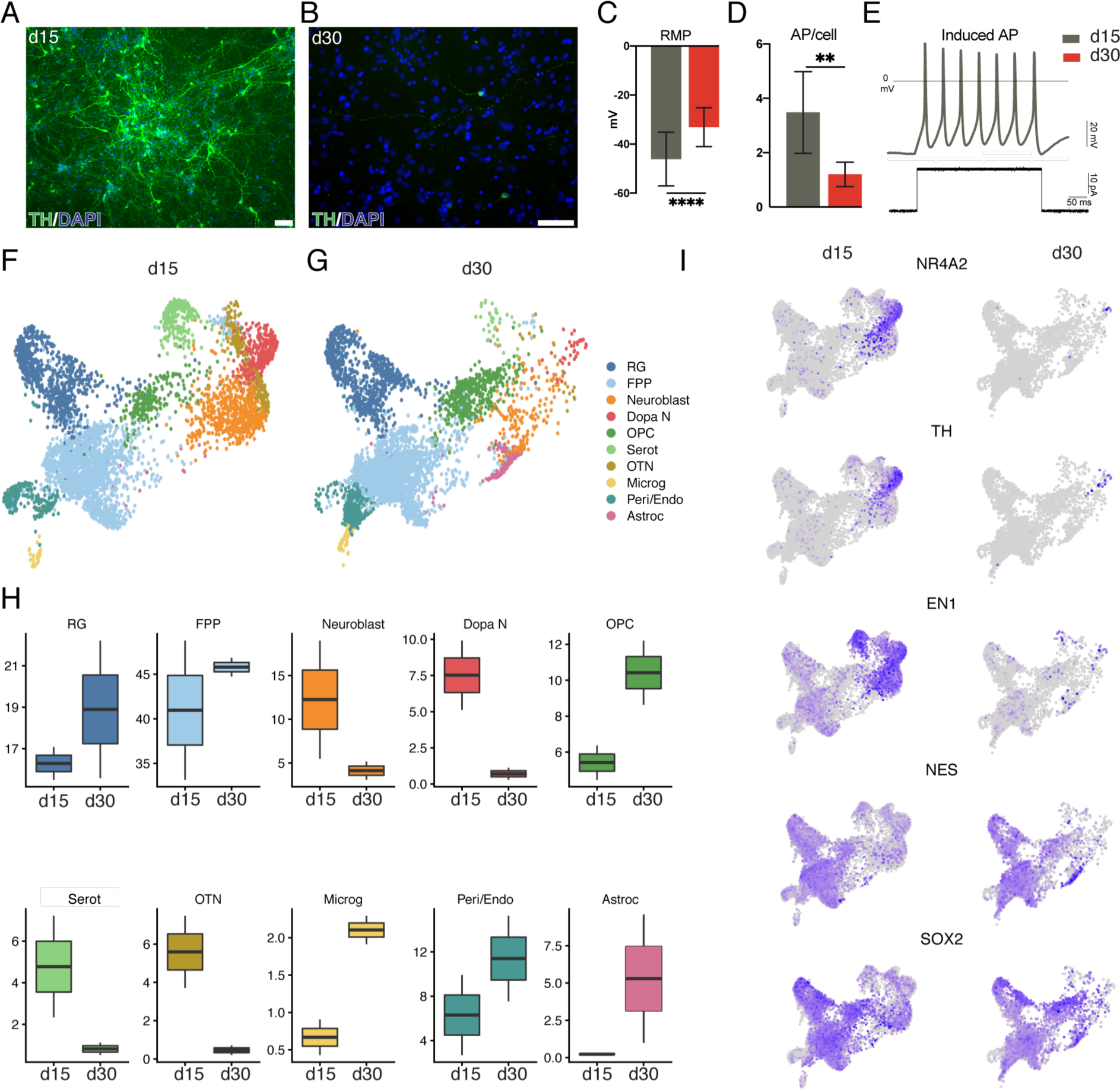
2D culture condition affects DA neuron maturation long term. **a**, Immunohistochemistry of TH in 2D culture at day 15 and **b**, at day 30. Scale bars, 100µM. **c**, Measurements from whole-cell patch-clamp recordings showing Resting Membrane Potential (RMP) at d15 (n=19) indicating the presence of neuronal cells within the cultures. Recordings were performed on 3 different human embryos for each time point. **d**, Measurements from whole-cell patch-clamp recordings showing mean of Action Potentials, APs, per cell at d15 and at day 30. **e**, Representative trace of multiple induced APs showing a neuronal profile of the patched cell. **f**, UMAP plot showing 2D cultured cells at day 15 and **g**, at day 30. **h**, Boxplot of percentage of cells belonging to each identified cluster at day 15 and day 30 from three independent embryos. **i**, Feature plots visualizing early and late DA markers across clusters at day 15 and day 30. Colors indicated expression level.

By comparing single cell datasets from these cultures at different timepoints (day 15 and day 30), we reconstructed the developmental framework of different cell types. scRNA-seq transcriptomics revealed a significant reduction in the entire neuronal compartment over time, including Serot (*SLC64a, GRIA4*) and OTN (*ISL1, PHOX2B*) clusters (Fig. 2H). In particular, the DA cluster was almost completely absent in 2D culture after only 1 month, preventing the recapitulation of more advanced and functional DA neurons (Fig. 2H,I). By contrast, RG and FP populations were preserved over time, while oligodendrocyte progenitors and astrocyte populations, which were almost absent at day 15, became clearly distinguishable as growing populations at 1 month (Fig. 2H).

### 3D culture recapitulates functionally mature human DA neurons

We hypothesized that fetal hVM cells cultured in 3D organoid-like structures could better maintain fetal DA neurons in long-term cultures. We therefore induced self-aggregation of 70,000 fetal VM cells using low attachment U-bottom plates (Fig. 3A). This gave rise to a cluster assembly by day 3, followed by complete 3D structure formation at day 15 (Fig. 3B). To provide structural support for these long-term cultures, drops of Matrigel were used to embed the clusters at day 30. Immunohistochemical analysis revealed the presence of TH neurons at day 15, similar to what we had observed in 2D primary cultures (Fig. 3C). However, in stark contrast to monolayer culture, the TH neurons in 3D structures were maintained at day 30 (Fig. 3D). Immunohistochemical analysis using antibodies for the mesodiencephalic FP/DA progenitor markers FOXA2 and OTX2 confirmed the midbrain identity of the cells (Fig. 3C and Supplementary Fig. 3A). At day 30, we also found the expression of mature DA markers such as DAT, dopa decarboxylase (AADC) and calcium-binding protein 1 (CALB1) (Fig. 3D and Supplementary Fig. 3B), indicating a mature subtype-specific identity of DA neurons in these cultures. Whole-cell patch-clamp recordings at day 30 revealed that the majority of cells (17/24) were able to fire multiple mature APs (Fig. 3F and Supplementary Fig. 3C). A third of the cells also showed spontaneous firing (8/24; Fig. 3G) and repetitive firing (8/24; Fig. 3H), a profile typical of mature DA neurons. The presence of postsynaptic activity was also detected in 10% of the cells (Fig. 3I). By performing both confocal spinning disk (Fig. 3E) imaging and iDISCO (Supplementary Fig. 3D and Supplementary Video 1), we obtained an anatomical 3D reconstruction of the complex DA neuronal circuitry and bundle connections within 3D fetal structures, showing that these DA neurons could be stably maintained for up to three months *in vitro*. To further assess whether this intricate DA network corresponded to a functional and active neuronal map, we performed calcium imaging of MAP2-GcaMP3-labeled neurons from 3D VM cultures (Fig. 3J and Supplementary Video 2) at 3 months. The presence of calcium waves with different kinetics in MAP2^+^ cells indicated active neuronal signaling (fig 3J), pointing to the fact that healthy and functionally mature neurons were present in these 3D fetal cultures.

**Figure 3.**
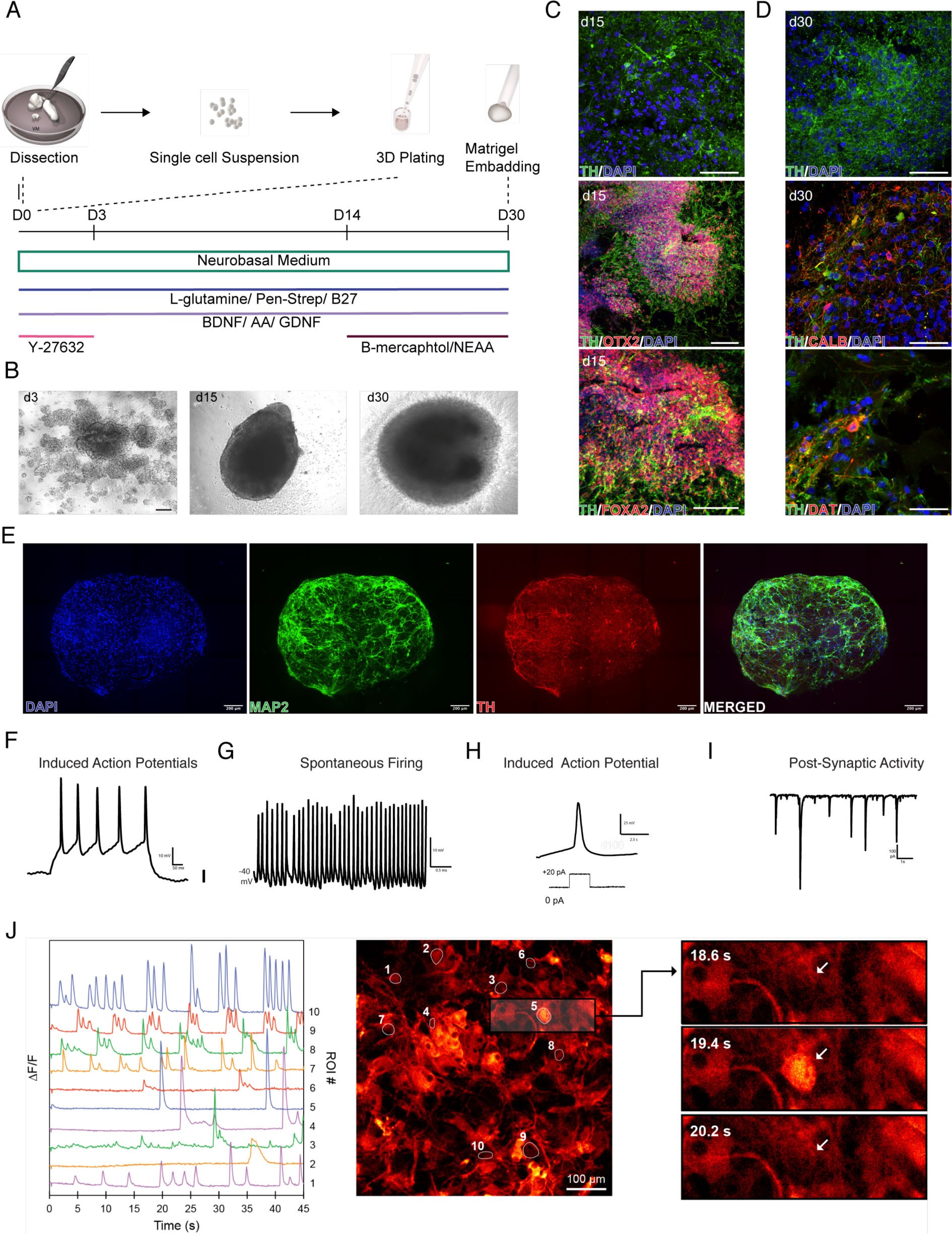
3D culture environment allows for the differentiation and maturation of human fetal DA. **a**, Schematic overview of protocol and experimental design. **b**, Representative bright-field images of 3D human fetal VM culture differentiation at different time points. Scale bar, 100 µM. **c**, Cryosection of 3D human fetal VM culture at day 15 showing TH/FOXA2 and TH/OTX2 double staining. Scale bars, 100 µM. **d**, Immunohistochemistry of TH/DAT and TH/CALB in 3D human fetal VM culture at day 30. Scale bars, 100 µM. **e**, 3D reconstruction of an image stack from 80 µm thick optical section of TH and MAP2 immunohistochemistry at day 120. Scale bars, 200 µM. **f**, Representative trace of Induced Action Potentials at d30 form 3D cultures indicative of mature neuronal profile. **g**, Representative trace of Spontaneous Firing indicative of a possible DA neuronal profile. **h**, Representative trace of Induced APs elicited by small step of current injection indicative of DA neuronal profile. **i**, Representative trace of Post-Synaptic activity indicative of an active neuronal network connection in the 3D system. **j**, Differential fluorescence intensity profile as a function of time for 3D hVM cultures at day 100 expressing MAP2-GCamP3 (left); fluorescence image with segmented regions of interest corresponding to individual cells (middle). Scale bar, 100 µM. Three timeframes displaying the change in intercellular fluorescence intensity for two cells indicated by arrows (right).

### scRNA-seq identifies cell diversity and distinct DA molecular identities in 3D human fetal VM organoid like cultures

Histological and functional analysis showed that DA neurons differentiated over time and that mature DA neurons were present in the long-term 3D cultures of human fetal VM (Fig. 3C,D,E and Supplementary Fig. 3A,B, D). To comprehensively characterize the cellular composition of these 3D cultures, we performed scRNA-seq at day 15 and day 30 (Fig. 4A,B). scRNA-seq confirmed that while the relative proportion of Serot and OTN neuron populations was decreasing over time in this 3D model, the DA neuron cluster was preserved and became the predominant neuronal cell type (Fig. 4A-C). Feature plots of well-established markers of early and late DA neurogenesis clearly visualized a robust DA population (Fig. 4D). While RG and FP populations did not vary significantly over time, microglia were almost absent after 15 days in 3D culture (Fig. 4A-C). Moreover, while oligodendrocyte progenitors emerged as a distinct cluster, astrocytes, previously captured as an expanding population in 2D culture, were almost completely absent at both timepoints (Fig. 4A-C).

**Figure 4.**
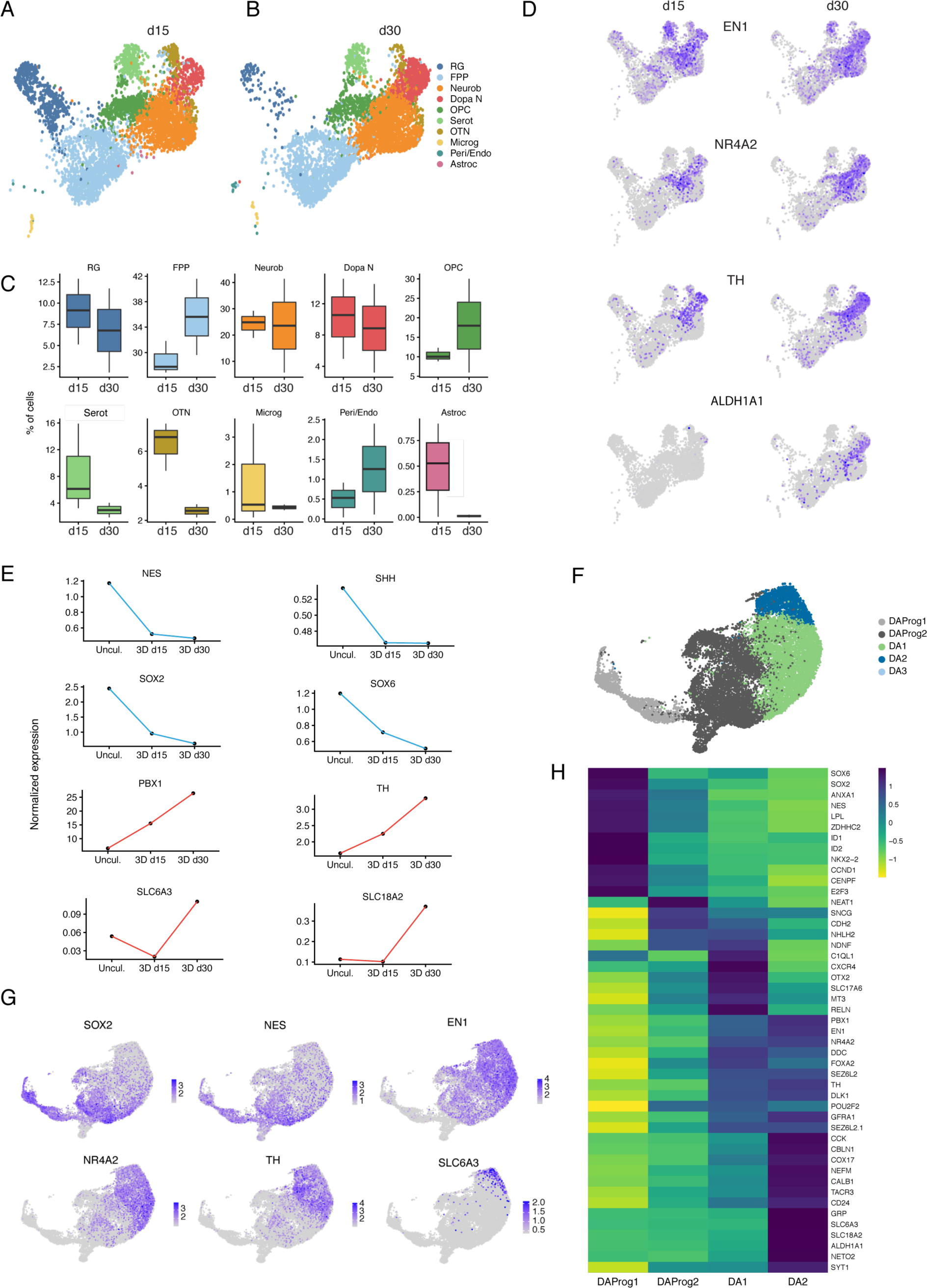
scRNA-seq captures molecular diversity of DA neuron within 3D fetal hVM culture. **a**, UMAP plot showing 3D human fetal VM culture at day 15 and **b**, at day 30. **c**, Boxplot of percentage of cells belonging to each identified cluster at day 15 and day 30 from three independent experiments. **d**, Feature plots visualizing early and late DA markers across clusters at day 15 and day 30. Colors indicate expression level. **e**, Pseudotemporal expression pattern of early and late DA markers from hVM fetal dissection after day15 and day30. **f**, UMAP plot showing DA sub-clusters from scRNA-seq dataset of 3D human fetal VM culture (day 15-30). **g**, Feature plots visualizing early and late DA markers in DA sub-clusters of 3D human fetal VM culture (day 15-30). **h**, Heatmap showing differentially expressed genes and manually selected markers in 4 DA neuron subclusters (DA- Prog1, DA- Prog2, DA-1 and DA-2).

We next pseudotemporally reconstructed the expression of the early and late DA markers that we had detected in the fetal VM tissue (Fig. 1I) and in the 3D cultures. The integrated scRNA-seq data showed that expression levels of late and mature DA markers such as *PBX1, TH, DAT* and *VMAT-2* increased over time with a corresponding decrease in *NES, SHH, SOX2* and *SOX6* (Fig. 4E). Overall, these findings validate our 3D culture system as a functional model able to mimic later stages of fetal DA neuron development in a dish.

The large number of functionally mature DA neurons present (n=12600) in fetal 3D culture enabled us to undertake a high-resolution analysis of DA neuron diversity at single cell level. We employed UMAP graph-based clustering to subcluster the DA population, which was found to be further segregated into four distinct major molecular identities (Fig. 4F). FeaturePlot analysis revealed that two populations were immature and shared molecular features of DA progenitors (*NES, SOX2, SOX6*). We identified these clusters, termed DAprog-1 and DAprog-2 (light and dark grey, respectively) as immature DA neurons not yet capable of neurotransmission (Fig. 4G). The other two clusters were more mature populations expressing high levels of late DA markers including *NR4A2, DDC*, and *TH*, here referred as DA-1 and DA-2 (Fig. 4G). Within DAprog-1 we found a proliferative signature (*CCND1, CENPF, E2F3*) together with pro-neural basic helix-loop-helix factor *ASCL1* expression (Fig. 4H). The neural stem/progenitor transcription factor *SOX2*, the PD neuroprotective factor *ANXA1*, and *SOX6*, known to have a crucial role in regulating specification of DA neurons in the substantia nigra [21], were also expressed (Fig. 4H). DAprog-2 was instead characterized by the loss of proliferative markers and pro-neural factors and acquired transcription factors regulating DA neuron specification such as *EN1, NURR1*, and *PBX1*. Interestingly, *SNCG*, encoding member of the synuclein family of proteins and the neural receptor *NETO2*, newly found in a human DA neuronal dataset, were also significantly seen in this subcluster.

Both DA-1 and DA-2 showed comparable enrichment in *TH* as well as in *DLK1* expression (Fig. 4H). However, DA-1 displayed lower levels of *DAT* and were primarily enriched for *OTX2, SLC17A6*, also known as *V-GLUT2*, and in *POU2F2*, which have recently been found in both human and mouse DA neuron scRNA-seq datasets (Fig. 4H) [9, 10]. DA-1 was molecularly defined also by expression of *C1ql*, which is involved in DA synapse formation and *CXCR4*, which is required for the migration and projection orientation of A9-A10 DA neurons. In contrast, DA-2 expressed high levels of *DAT* and was also enriched in the enzyme involved in DA catabolism and transport (*ALDH1a1, VMAT2*), as well as in *PITX3*, together with its transcriptional co-activator LMO3 (Fig. 4H).

## CONCLUSIONS

Animal models have been instrumental in understanding neurodevelopmental and neurodegenerative disorders, but their limitations in revealing key features of developmental, genetic, and pathological mechanisms unique to humans are increasingly recognized. However, the inaccessibility of human brain tissue makes such studies extremely challenging, and PSC-based models do not capture all aspects of human development. Here, we performed extensive scRNA-seq analysis to decode VM development in human embryos from onset to peak DA-genesis and also profiled functionally mature DA neurons in 3D cultures derived from the human fetal VM.

Two previous studies [9, 10] reported similar transcriptional profiling of fetal VM using smartseq2. In this study, we used high-throughput droplet-based seq (10x) to enable analysis of much larger cell numbers and obtained data from a total of 23,483 VM cells from three fetuses of gestational age 6–11 weeks PC, which is considerably greater than in previously published datasets [4, 10]. The cells separated into 10 distinct clusters. As expected, the majority of the cells at these early developmental timepoints were RG and FPP, in agreement to what was observed in the study by La Manno et al. However, three distinct neuronal subtypes were also detected: DA neurons, OTN neurons, which are formed in VM, and a small population of serotoninergic neurons, known to be located in close proximity to DA neurons during development [22]. Moreover, already at these early developmental stages, DA neurons/progenitors of molecularly distinct identities were present, suggesting an early emergence of molecular sub-identities.

Given the even more limited accessibility of fetal tissue after the first trimester and the inherent problems with using such tissue, we established primary VM cultures in order to profile the functionally mature human DA neurons. We first performed traditional monolayer (2D) cultures but this proved inadequate so we then developed a 3D organoid like culture system that allowed for the long term maturation of these dopaminergic cells [23, 24],. These 3D culture allowed us therefore to study the developmental trajectory of DA progenitors to functionally mature DA neurons, where they were found to exhibit high expression of TH as well as enzymatic and transporter components of DA neurotransmission.

This new culture system enabled targeted transcriptional analysis of 2500 human midbrain DA neurons at a stage where they have reached functional maturation. This analysis revealed four major populations: two were characterized by expression of pro-neural factors and markers of DA progenitors, whereas the expression of *TH* and components of neurotransmission defined two more mature DA populations. A previous study sequencing fetal VM tissue using smartseq2 also identified distinct DA subclusters named DA0, DA1 and DA2 (La Manno et al., Cell, 2017). In line with our data, these subclusters differed based on expression of genes such as *TH, PITX3, EN1*, and *TMCC3* (DA0, DA1, DA2), *SLC6A3, NETO2*, and *KCNJ6* (DA1, DA2), and *LMO3* and *ALDH1A1* (DA2). Most of these genes were also detected in our dataset but did not always segregate into the same clusters. This may partly be due to differences in the sequencing methods adopted (smartseq2 vs 10x) and cell numbers analyzed (122 analyzed in La Manno et al., and 2776 in this study), but more likely reflects that the molecular subtype identities are refined as the DA neurons mature in culture.

Transcriptional profiling at the single cell level of the adult human midbrain will be of great value in understanding how molecular distinct subtypes related to the classical definition of A9 and A10 neurons arise. Of note, the expression of Girk2 and Calb, commonly used to segregate the A9 and A10 DA neurons in the adult midbrain, was not found to be clearly enriched in our different DA subclusters nor in the Le Manno data sets, suggesting that at this developmental stage they had not yet segregated into specific DA neuron subtypes. However, *ALDH1a1* was expressed specifically in cluster DA2, suggesting its expression may be an earlier marker to identify subtype-specific DA neurons.

Comparative studies of fetal ventral midbrain (VM)-derived DA neurons and stem cell-derived DA neurons and their progenitors in terms of gene expression, phenotypic identity, and functional properties after transplantation have been key in establishing differentiation protocols and advancing stem cell derived DA neurons towards clinical use [4–7]. To more precisely control DA subtype identity in stem cell cultures, it is also essential to map their molecular diversity during development and in the adult brain. The data presented in this study provide a unique molecular characterization of developing and functionally mature human DA neurons and will serve as a valuable resource for future advancements using stem cell-derived and reprogrammed human DA neurons *in vitro* and *in vivo*.

## EXPERIMENTAL PROCEDURES

### Human embryonic tissue source

Human fetal tissues were collected from 6–11 week PC legally terminated embryos at Malmö Hospital (Malmö, Sweden) and Addenbrooke’s Hospital (Cambridge, U.K.). Ethical approval for the use of postmortem human fetal tissue was provided by the Swedish National Board of Health and Welfare in accordance with existing guidelines, including informed consent from women seeking abortions, and by the National Research Ethics Service Committee East of England - Cambridge Central (Local Research Ethics Committee, reference no. 96/085). The gestational age of each embryo was determined by crown-to-rump length (CRL) measured at either the time of dissection when the quality of the embryo allowed for this or otherwise estimated by ultrasound measurements prior to abortion. The external features of the embryo were also carefully monitored to confirm that the CRL correlated with the appropriate embryonic stage. Samples from U.K. were shipped overnight on ice in HIBERNATE media to Sweden.

### Acutely dissociated cell preparations and culture conditions

Tissue from both Sweden and U.K. was dissected in HIBERNATE media. Narrow sub-dissection of the human VM was performed and the tissue washed in phosphate buffered saline (PBS) solution. After 3 washes, the tissue was treated with Accutase (PAA Laboratories) for 20 min at 37°C degrees. After incubation, single cell suspensions were generated by mechanical dissociation and the cells plated at a density of 70,000 cells/well (36,842 cells/cm^2^) in culture media. Culture media used was formulated as follows: Neurobasal Medium, 2 nM L-glutamine, 100 µg/mL pen/strep, 20 ng/ml BDNF, 10 ng/ml GDNF, 0.2 mM AA, 1/3 B27. On the plating day after dissociation, the culture media was supplemented with Y-27632 (10 µM) to improve neuronal survival. A total of 1% minimum essential medium-non essential amino acids (MEM-NEAA) and 0.1% 2-mercaptoethanol was added to the culture media from day 14. Media was changed every 2 days. 2D cultures were performed in standard plates coated with a combination of polyornithine (15 µg/mL), fibronectin (0.5 ng/µL) and laminin (5 µg/mL). 3D cultures were generated using U-bottom shaped ultra-low attachment 96-well plates (Corning). Droplets of Matrigel were applied to allow embedding at day 30 to sustain long-term cultures. At the time of embedding, 3D hVM cultures were transferred into ultra-low attachment 24-well plates (Corning). 3D cultured organoids used for calcium imaging were left attached on glass coverslips coated with polyornithine, fibronectin, and laminin at day 90.

### 3D hVM culture preparation for GCaMP3 recordings

At day 95, when the 3D hVM cultures had attached onto the glass coverslips, cells were transduced with the lentivirus MAP2-GCaMP3 in culture media overnight. The following day, a complete media change was performed. Prior to calcium imaging recordings, media was changed regularly every 2 days.

### Immunocytochemistry and immunohistochemistry

The cells were fixed in 4% paraformaldehyde solution for 15 min at room temperature (RT) prior to staining. The cells were pre-incubated in a blocking solution containing 0.1 M PBS with potassium (KPBS) + 0.1% Triton + 5% serum (of secondary antibody host species) for 1– 3 hours before the primary antibody solution was added.

3D cultures were fixed in 4% paraformaldehyde solution overnight at RT and cryoprotected in 30% sucrose before being frozen in Tissue-Tek O.C.T (Sakura FineTek, Europe BF). Sectioning of the frozen 3D samples was performed using a cryostat supplier, with slices of 200 um thickness.

The cells were incubated with the primary antibodies overnight at 4°C and the following day they were washed with KPBS before adding the secondary antibody solution containing fluorophore-conjugated antibodies (1:200, Jackson ImmunoResearch Laboratories) and DAPI (1:500). The cells were incubated with the secondary antibodies for 2 hours at RT and finally washed with KPBS.

Primary antibodies used were: rabbit anti-tyrosine hydroxylase (TH) (1:1,000, Pel-Freeze Biological), mouse anti-SOX2 (1:50, R&D Systems), guinea pig anti-GLAST (1:1,000, Chemicon), mouse anti-nestin (1:200 *in vivo*, 1:500 *in vitro*, BD Bioscience), chicken anti-vimentin (1:10,000, Chemicon), goat anti-vimentin (1:1,000 *in vitro*, Sigma), goat anti-neurogenin2 (Ngn2, 1:500, Santa Cruz), mouse anti-microtubule-associated protein 2 (Map2, 1:250, Sigma), rabbit anti-glial fibrillary acidic protein (GFAP, 1:1,000, DAKO), mouse anti-4A4 (1:2,000, Medical & Biological Laboratories), rabbit anti-BLBP (1:5,000, Chemicon), rabbit anti-PAX6 (1:500, BioSite) and rabbit anti-LMX1a (1:10,000, kindly donated by Dr German).

### scRNA-seq analysis

Cell suspensions were loaded into a 10x Genomics Chromium Single Cell System (10x Genomics) and libraries were generated using version 3 chemistry according to the manufacturer’s instructions. Libraries were sequenced on Illumina NextSeq500 (400 million reads) using the recommended read length. Sequencing data was first pre-processed through the Cell Ranger pipeline (10x Genomics, Cellranger count v2) with default parameters (expect-cells set to number of cells added to 10x system), aligned to GrCH38 (v3.1.0) and resulting matrix files were used for subsequent bioinformatic analysis. Seurat (version 3.1.1 and R version 3.6.1) was utilized for downstream analysis. Batch effects were removed using the Harmony algorithm (1.0), treating individual 10x runs as a batch. Cells with at least 200 detected genes were retained and the data was normalized to transcript copies per 10,000, and log-normalized to reduce sequencing depth variability. For visualization and clustering, manifolds were calculated using UMAP methods (*RunUMAP*, Seurat) and 20 precomputed principal components and the shared nearest neighbor algorithm modularity optimization based clustering algorithm (FindClusters, Seurat) with a resolution of 0.2. Identification of differentially expressed genes between clusters was carried out using the default Wilcoxon rank sum test (Seurat) and gene ontology overrepresentation analysis was performed using the enrichGO function in the clusterProfiler package (3.13) using MSigDB as the database.

### iDISCO

3D fetal structures were fixed in 2% paraformaldehyde overnight at 4°C followed by permeabilization in 0.2% Triton X-100/20% DMSO. After 2 hours, organoids were incubated overnight in 0.1% Triton X-100, 0.1%Tween20, 0.1% C_24_H_39_NA0_4_, 0.1% NP40, and 20% DMSO at 37°C.

Cultures were incubated with primary antibodies for 2 days at 37°C followed by 2 days’ incubation with secondary antibodies with embedding in 1% agarose and dehydrated in ascending concentration series of methanol and dichloromethane as previously described [25].

### Microscopy

Fluorescent images were captured using a Leica DMI6000B widefield microscope. Image acquisition software was Leica LAS X and images were processed using Adobe Photoshop CC 2018. Any adjustments were applied equally across the entire image, and without the loss of any information.

Immunohistochemical stainings were analyzed by confocal microscope (Leica) at 20× or 63× resolution. Double staining was confirmed by conduction of high-magnified confocal Z-stacks. All figures were assembled in Canvas software.

### Calcium imaging of MAP2-GCamP3-labeled neurons

Calcium imaging was performed at day 100 of 3D hVM cultures containing the MAP2-GCamP3 reporter. Cell culture media was replaced with 100 µL baseline buffer containing 1.2 × 10^−3^ M MgCl_2_, 2 × 10^−3^ M CaCl_2_, 150 × 10^−3^ M NaCl, 5 × 10^−3^ M KCl, 5 × 10^−3^ M glucose, and 10 × 10^−3^ M HEPES. Imaging was performed on an inverted Ti2 microscope (Nikon) equipped with a CSU-W1 spinning disc system (Yokogawa), a sCMOS camera (Teledyne Photometrics), and a 20× objective. An environment control chamber was used to maintain the temperature at 37°C and CO_2_ level at 5% during imaging. Exposure time was set to 30 ms or 100 ms depending on the dynamics of calcium transients. Spontaneous activity was recorded from 3 different 3D fetal structures from different embryos. Images were analyzed in ImageJ (NIH) and plotted in QtiPlot.

### Electrophysiology

Whole-cell patch-clamp electrophysiological recordings were performed at day 15 for 2D and day 30–100 for 3D hVM cultures. Cells from the 2D condition model were cultured on glass coverslips and transferred to a recording chamber with constant flow of Krebs solution gassed with 95% O_2_-5% CO_2_ at RT. The composition of the standard solution was (in mM): 119 NaCl, 2.5 KCl, 1.3 MgSO_4_, 2.5 CaCl_2_, 25 glucose, and 26 NaHCO_3_. 3D cultured hVMs were maintained in ultra-low attachment plates, and on recording day were transferred to a recording chamber with Krebs solution gassed with 95% O_2_-5% CO_2_ at RT that was manually exchanged at the end of each individual cell recording. For recordings, Multiclamp 700B amplifier (Molecular Devices) was used together with borosilicate glass pipettes (3–7 MOhm) filled with the following intracellular solution (in mM): 122.5 C6H11KO7, 12.5 KCl, 0.2 EGTA, 10 HEPES, 2 MgATP, 0.3 Na_3_GTP, and 8 NaCl adjusted to pH 7.3 with KOH as previously described [26]. Data acquisition was performed with pClamp 10.2 (Molecular Devices); current was filtered at 0.1 kHz and digitized at 2kHz. Cells with a neuronal morphology and round cell body were selected for recordings. RMPs were monitored immediately after breaking-in in current-clamp mode. Thereafter, cells were kept at a membrane potential of −45 mV to −70mV, and 500 ms currents were injected from −50 pA to +100 pA with 10 pA increments to induce action potentials. For inward sodium and delayed rectifying potassium current measurements, cells were clamped at −70mV and voltage-depolarizing steps were delivered for 100 ms at 10 mV increments. Spontaneous APs were recorded in current-clamp mode at RMPs. Post-synaptic activity was recorded in voltage-clamp mode at RMPs.

### Statistical analysis

All data is expressed as mean +/-standard error of the mean (SEM). A Shapiro-Wilk normality test was used to assess the normality of the distribution and parametric or nonparametric tests were performed accordingly. Statistical analyses were conducted using GraphPad Prism 8.0.

## AUTHOR CONTRIBUTION

M.P., M.B., A.F. conceived the project, designed and interpreted data and analyzed results; M.B., A.F., J.K., E.S., F.N., B.M and DRO performed experiments, interpreted data and analyzed results; P.S. and Y.S. performed bioinformatic analysis; J.N.W. performed tissue preparation for fetal experiments; R.B. designed experiments and interpreted results;. M.P., M.B. and AF wrote the paper with input from all authors.

## ACKNOWLEDGEMENTS

We thank Sol Da Rocha Baez, Jenny Johansson, Ulla Jarl and Marie Persson Vejgården for invaluable technical assistance.

The research leading to these results has received funding from the New York Stem Cell Foundation (MP), the European Research Council (ERC Grant Agreement no. 771427, MP), the Swedish Research Council (grant agreements 2016-00873 [MP]), the Swedish Parkinson Foundation (Parkinsonfonden, MP), the Swedish Brain Foundation (Hjärnfonden FO2019-0301, MP), the Strategic Research Area at Lund University MultiPark (Multidisciplinary research in Parkinson’s disease) and KAW (2018.0040). M. Birtele is supported by a BMT Marie Curie Horizon 2020 fellowship. M. Parmar is a New York Stem Cell Foundation– Robertson Investigator.

---

**supplementary Fig 1.**
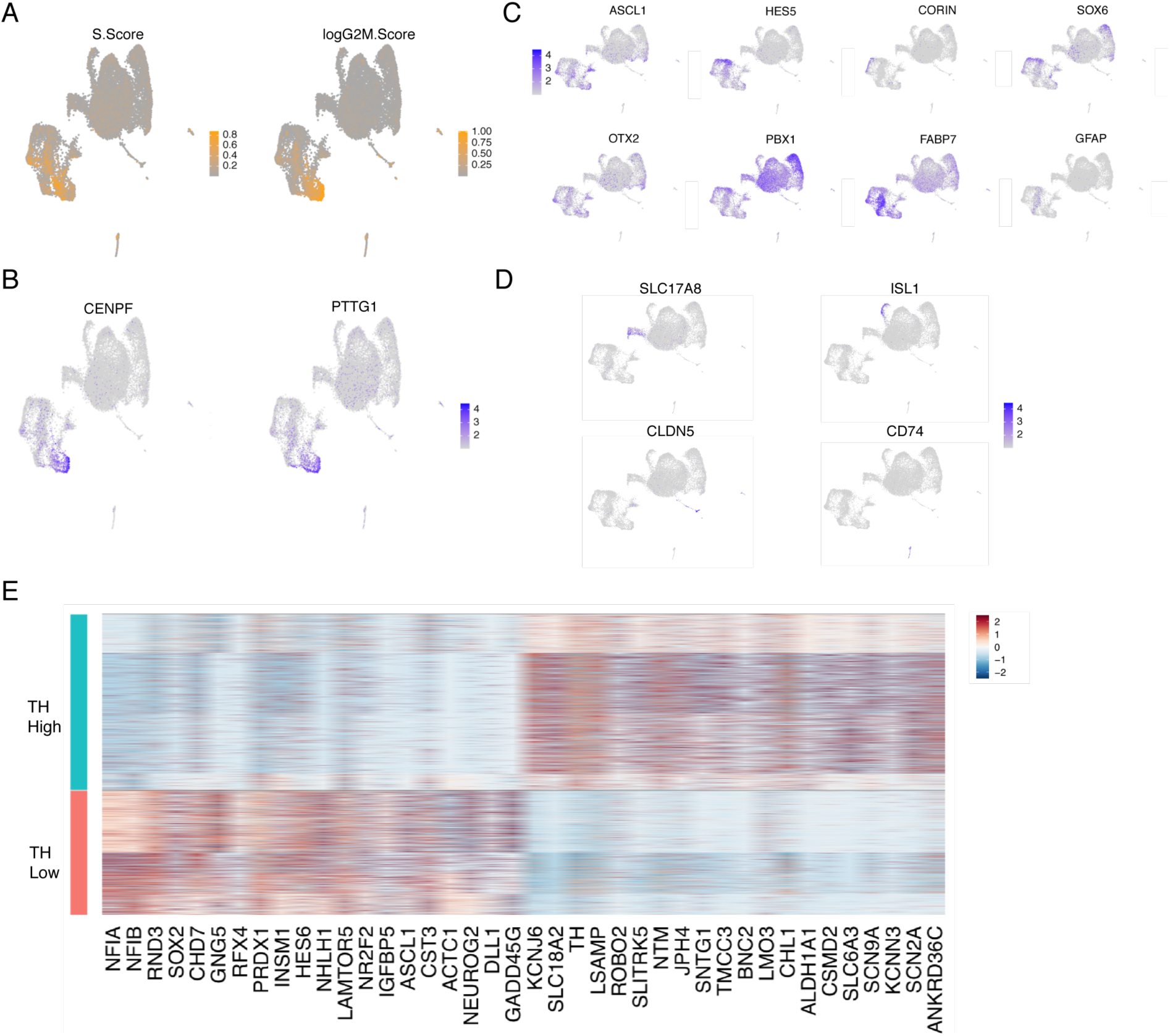

**suppl Fig 2.**
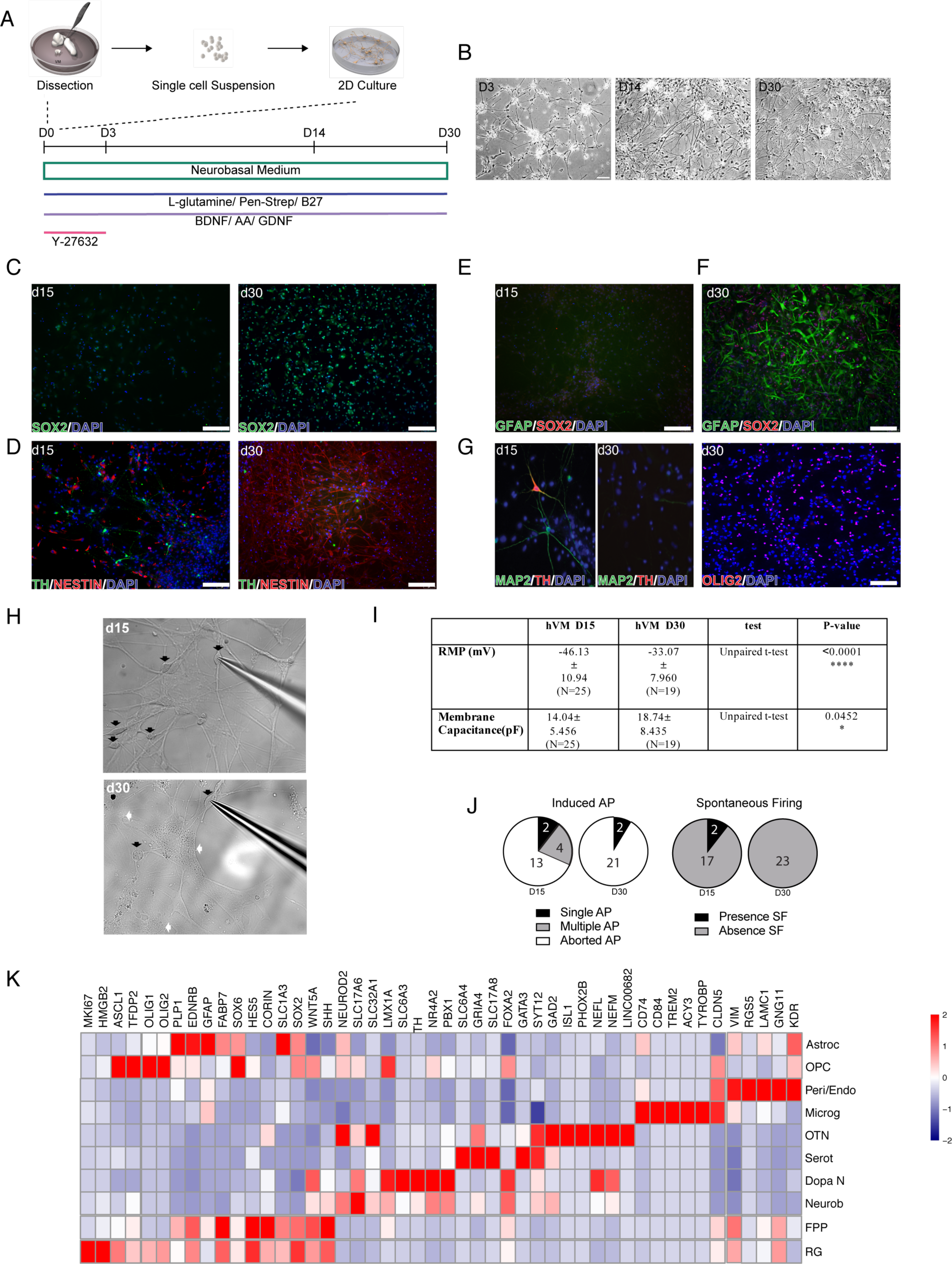

**suppl Fig 3.**
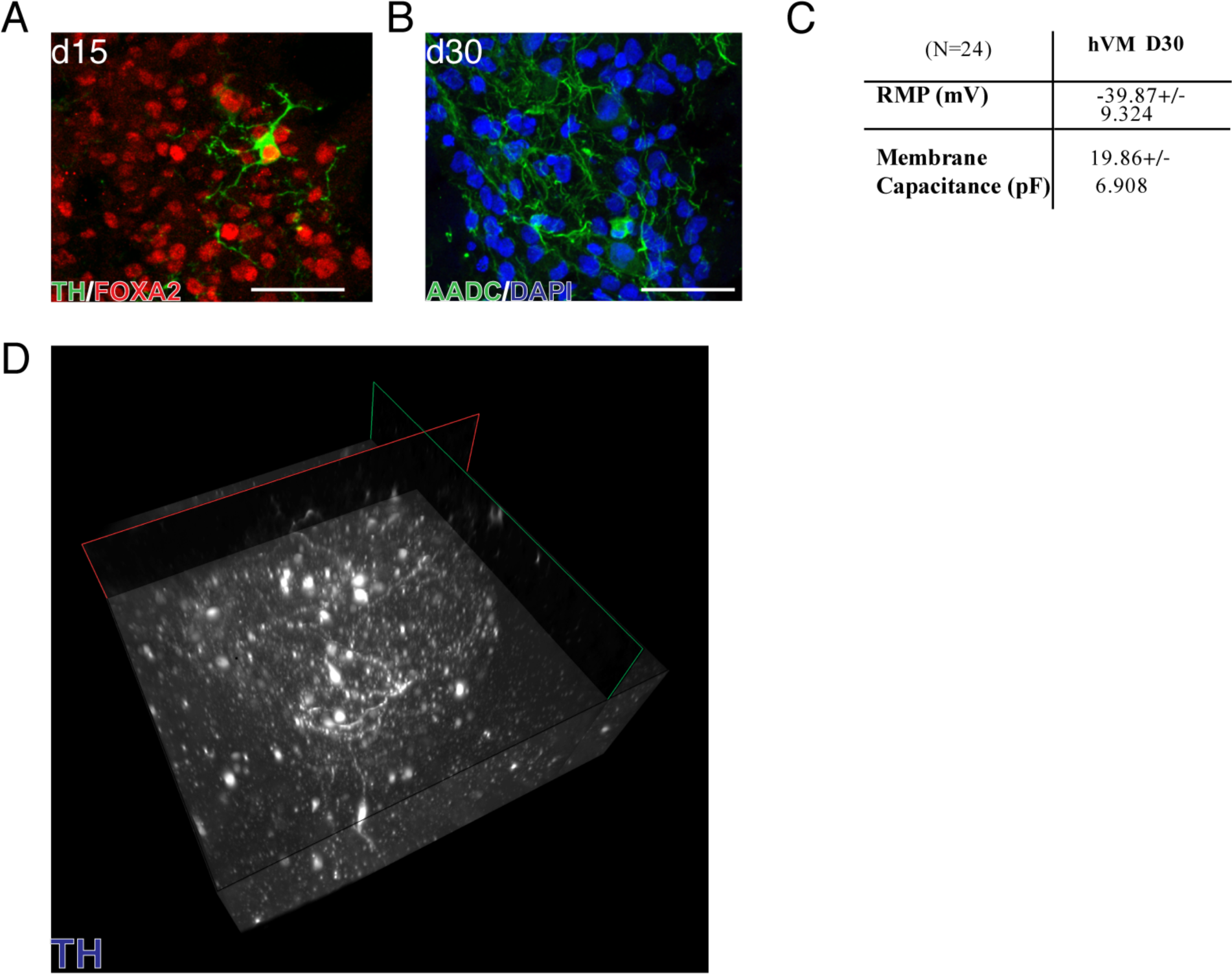

## Notes

### Competing Interest Statement

The authors have declared no competing interest.

## References

1. Barker RA, Farrell K, Guzman NV, He X, Lazic SE, Moore S, Morris R, Tyers P, Wijeyekoon R, Daft D, Hewitt S, Dayal V, Foltynie T, Kefalopoulou Z, Mahlknecht P, Lao-Kaim NP, Piccini P, Bjartmarz H, Björklund A, Lindvall O, Nelander-Wahlestedt J, Parmar M, Paul G, Widner H, Church A, Dunnett S, Peall K, Rosser A, Gurruchaga JM, Palfi S, Piroth T, Winkler C (2019) Designing stem-cell-based dopamine cell replacement trials for Parkinson’s disease. Nature Medicine 25:. doi: 10.1038/s41591-019-0507-2

2. Barker RA, Drouin-Ouellet J, Parmar M (2015) Cell-based therapies for Parkinson disease-past insights and future potential. Nature Reviews Neurology 11:492–503

3. Barker RA, Parmar M, Studer L, Takahashi J (2017) Human Trials of Stem Cell-Derived Dopamine Neurons for Parkinson’s Disease: Dawn of a New Era. Cell Stem Cell 21:569–573

4. Tiklová K, Nolbrant S, Fiorenzano A, Björklund ÅK, Sharma Y, Heuer A, Gillberg L, Hoban DB, Cardoso T, Adler AF, Birtele M, Lundén-Miguel H, Volakakis N, Kirkeby A, Perlmann T, Parmar M (2020) Single cell transcriptomics identifies stem cell-derived graft composition in a model of Parkinson’s disease. Nature Communications 11:. doi: 10.1038/s41467-020-16225-5

5. Kikuchi T, Morizane A, Doi D, Magotani H, Onoe H, Hayashi T, Mizuma H, Takara S, Takahashi R, Inoue H, Morita S, Yamamoto M, Okita K, Nakagawa M, Parmar M, Takahashi J (2017) Human iPS cell-derived dopaminergic neurons function in a primate Parkinson’s disease model. Nature 548:592–596. doi: 10.1038/nature23664

6. Grealish S, Diguet E, Kirkeby A, Mattsson B, Heuer A, Bramoulle Y, Van Camp N, Perrier AL, Hantraye P, Björklund A, Parmar M (2014) Human ESC-derived dopamine neurons show similar preclinical efficacy and potency to fetal neurons when grafted in a rat model of Parkinson’s disease. Cell Stem Cell 15:653–665. doi: 10.1016/j.stem.2014.09.017

7. Kirkeby A, Grealish S, Wolf DA, Nelander J, Wood J, Lundblad M, Lindvall O, Parmar M (2012) Generation of Regionally Specified Neural Progenitors and Functional Neurons from Human Embryonic Stem Cells under Defined Conditions. Cell Reports 1:703–714. doi: 10.1016/j.celrep.2012.04.009

8. Björklund A, Dunnett SB (2007) Dopamine neuron systems in the brain: an update. Trends in Neurosciences 30:194–202

9. Tiklová K, Björklund ÅK, Lahti L, Fiorenzano A, Nolbrant S, Gillberg L, Volakakis N, Yokota C, Hilscher MM, Hauling T, Holmström F, Joodmardi E, Nilsson M, Parmar M, Perlmann T (2019) Single-cell RNA sequencing reveals midbrain dopamine neuron diversity emerging during mouse brain development. Nature Communications 10:1–12. doi: 10.1038/s41467-019-08453-1

10. La Manno G, Gyllborg D, Codeluppi S, Nishimura K, Salto C, Zeisel A, Borm LE, Stott SRW, Toledo EM, Villaescusa JC, Lönnerberg P, Ryge J, Barker RA, Arenas E, Linnarsson S (2016) Molecular Diversity of Midbrain Development in Mouse, Human, and Stem Cells. Cell 167:566-580.e19. doi: 10.1016/j.cell.2016.09.027

11. Poulin JF, Zou J, Drouin-Ouellet J, Kim KYA, Cicchetti F, Awatramani RB (2014) Defining midbrain dopaminergic neuron diversity by single-cell gene expression profiling. Cell Reports 9:930–943. doi: 10.1016/j.celrep.2014.10.008

12. Rosenberg AB, Roco CM, Muscat RA, Kuchina A, Sample P, Yao Z, Graybuck LT, Peeler DJ, Mukherjee S, Chen W, Pun SH, Sellers DL, Tasic B, Seelig G (2018) Single-cell profiling of the developing mouse brain and spinal cord with split-pool barcoding. Science 360:176–182. doi: 10.1126/science.aam8999

13. Cao J, Packer JS, Ramani V, Cusanovich DA, Huynh C, Daza R, Qiu X, Lee C, Furlan SN, Steemers FJ, Adey A, Waterston RH, Trapnell C, Shendure J (2017) Comprehensive single-cell transcriptional profiling of a multicellular organism. Science 357:661–667. doi: 10.1126/science.aam8940

14. Saunders A, Macosko EZ, Wysoker A, Goldman M, Krienen FM, de Rivera H, Bien E, Baum M, Bortolin L, Wang S, Goeva A, Nemesh J, Kamitaki N, Brumbaugh S, Kulp D, McCarroll SA (2018) Molecular Diversity and Specializations among the Cells of the Adult Mouse Brain. Cell 174:1015-1030.e16. doi: 10.1016/j.cell.2018.07.028

15. Kee N, Volakakis N, Kirkeby A, Dahl L, Storvall H, Nolbrant S, Lahti L, Björklund ÅK, Gillberg L, Joodmardi E, Sandberg R, Parmar M, Perlmann T (2017) Single-Cell Analysis Reveals a Close Relationship between Differentiating Dopamine and Subthalamic Nucleus Neuronal Lineages. Cell Stem Cell 20:29–40. doi: 10.1016/j.stem.2016.10.003

16. Hook PW, McClymont SA, Cannon GH, Law WD, Morton AJ, Goff LA, McCallion AS (2018) Single-Cell RNA-Seq of Mouse Dopaminergic Neurons Informs Candidate Gene Selection for Sporadic Parkinson Disease. American Journal of Human Genetics 102:427–446. doi: 10.1016/j.ajhg.2018.02.001

17. Arenas E, Denham M, Villaescusa JC (2015) How to make a midbrain dopaminergic neuron. Development 142:1918–1936. doi: 10.1242/dev.097394

18. Nelander J, Hebsgaard JB, Parmar M (2009) Organization of the human embryonic ventral mesencephalon. Gene Expression Patterns 9:555–561. doi: 10.1016/j.gep.2009.10.002

19. Almqvist PM, Åkesson E, Wahlberg LU, Pschera H, Seiger Å, Sundström E (1996) First trimester development of the human nigrostriatal dopamine system. Experimental Neurology 139:227–237. doi: 10.1006/exnr.1996.0096

20. Freeman TB, Spence MS, Boss BD, Spector DH, Strecker RE, Olanow CW, Kordower JH (1991) Development of dopaminergic neurons in the human substantia nigra. Experimental Neurology 113:344–353. doi: 10.1016/0014-4886(91)90025-8

21. Panman L, Papathanou M, Laguna A, Oosterveen T, Volakakis N, Acampora D, Kurtsdotter I, Yoshitake T, Kehr J, Joodmardi E, Muhr J, Simeone A, Ericson J, Perlmann T (2014) Sox6 and Otx2 control the specification of substantia nigra and ventral tegmental area dopamine neurons. Cell Reports 8:1018–1025. doi: 10.1016/j.celrep.2014.07.016

22. Marklund U, Alekseenko Z, Andersson E, Falci S, Westgren M, Perlmann T, Graham A, Sundström E, Ericson J (2014) Detailed expression analysis of regulatory genes in the early developing human neural tube. Stem Cells and Development 23:5–15. doi: 10.1089/scd.2013.0309

23. Lancaster MA, Knoblich JA (2014) Generation of cerebral organoids from human pluripotent stem cells. Nature Protocols 9:2329–2340. doi: 10.1038/nprot.2014.158

24. Quadrato G, Nguyen T, Macosko EZ, Sherwood JL, Yang SM, Berger DR, Maria N, Scholvin J, Goldman M, Kinney JP, Boyden ES, Lichtman JW, Williams ZM, McCarroll SA, Arlotta P (2017) Cell diversity and network dynamics in photosensitive human brain organoids. Nature 545:48–53. doi: 10.1038/nature22047

25. Renier N, Wu Z, Simon DJ, Yang J, Ariel P, Tessier-Lavigne M (2014) IDISCO: A simple, rapid method to immunolabel large tissue samples for volume imaging. Cell 159:896–910. doi: 10.1016/j.cell.2014.10.010

26. Pfisterer U, Kirkeby A, Torper O, Wood J, Nelander J, Dufour A, Björklund A, Lindvall O, Jakobsson J, Parmar M (2011) Direct conversion of human fibroblasts to dopaminergic neurons. Proceedings of the National Academy of Sciences of the United States of America 108:10343–10348. doi: 10.1073/pnas.1105135108

